# Human antibody heavy chain variable region exon interacts with 3’ regulatory region to form a stable somatic hypermutation center

**DOI:** 10.64898/2026.07.20.739632

**Authors:** Ying Guan, Mariia Mikhova, Zhuo Yang, Kai Xu, Shuxian Xie, Xin Lin, Adam Yongxin Ye, Taimor Williams, Jianshu Wang, David G. Schatz, Frederick W. Alt, Xi Chen

**Author notes:** Frederick W. Alt and Xi Chen. **Email:**. Y.G. and X.C. contributed equally to this work. Present address: University of California San Diego, La Jolla, CA 92093. This study is conducted in Frederick W. Alt lab. **Author Contributions:** Y.G., F.W.A., and X.C. designed research; Y.G., M.M., Z.Y., S.X., and X.C. performed research; K.X., X.L., A.Y.Y., and T.W. contributed new reagents/analytic tools; Y.G., F.W.A., and X.C. analyzed data; and Y.G., F.W.A., and X.C. wrote the paper. D.G.S. provided AID7.3in cells and D.G.S. and J.W. contributed to interpreting data and polishing the paper. **Competing Interest Statement:** The authors declare no competing interest.

## Abstract

Activation-induced cytidine deaminase (AID) initiates somatic hypermutation (SHM) of immunoglobulin heavy chain (HC) variable region exons in germinal center (GC) B cells, allowing selection of affinity-matured B cell receptors. The V(D)J exon location is ‘‘privileged’’ for SHM targeting of diverse sequences within it. In analogy to the related HC class switch recombination center, we proposed that cohesin-mediated loop extrusion juxtaposes widely separated enhancers, including 3’ *Igh* regulatory region (“3’RR”), with the V(D)J exon in GC B cells to establish a SHM-center (SHM-C) “privileged” for AID access. To test this hypothesis, we employed a human RAMOS GC B lymphoma cell SHM model with inducible AID expression in which we targeted an unmutated mouse HC V(D)J exon in place of its human counterpart. Activation of AID expression in this cycling RAMOS line for 10 days led to robust SHM accumulation in the mouse V(D)J exon with a pattern similar to that in mouse GC B cells. In this model, the mouse V(D)J exon was indeed juxtaposed to the two human 3’RRs and other elements consistent with SHM-C formation. While deletion of individual 3’RRs had little effect, deletion of both greatly reduced SHM, but left V(D)J exon transcription unabated. Moreover, the SHM-C and SHM accumulation remained intact over a 10-day period in viably G1-arrested and viably G1-arrested RAD21 (cohesin)-depleted versions of the model. We discuss implications for SHM-C structure and function, its long-term functional stability in the absence of the loop extrusion process proposed to assemble it, and for 3’RR SHM functions beyond transcriptional activation.

**Significance Statement:** Upon encountering pathogens, B cells enter specialized lymphoid structures known as germinal centers (GCs). GC B cells express activation-induced cytidine deaminase (AID) that initiates somatic hypermutation (SHM) of the portion of antibody genes that encodes antigen recognition. B cells in which mutations increase antibody affinity for activating pathogen are selected to enhance antibody responses. We proposed SHM occurs in a chromosomal structure in the antibody gene locus termed a SHM-center (SHM-C). By studying SHM in a human B cell lymphoma model, we provide direct evidence for existence of a SHM-C assembled from different functional components normally separated by considerable distances. We also find that the SHM-C is surprisingly stable and continues to function for many days in non-dividing cells.

## Introduction

The variable region exons of the immunoglobulin heavy chain (*Igh*) locus are assembled in developing B cells through V(D)J recombination. V(D)J recombination generates the primary B cell receptor repertoire (1). Upon exposure to antigen, B cells can undergo two further genomic modifications both initiated by the activation-induced cytidine deaminase (AID) (2). AID initiates *Igh* class switch recombination (CSR) in CSR-activated B cells to exchange expressed *Igh* constant region exons (C_H_s) (3). In germinal center (GC) B cells, AID also initiates antigen-induced *Igh* V(D)J exon somatic hypermutation (SHM), allowing affinity maturation of B cell receptors (BCRs) (4, 5). AID initiates both processes by deaminating cytosines within short target motifs that are converted by DNA repair pathways into DNA double-strand breaks (DSBs) for CSR and, predominantly, focal point mutations in V(D)J exons (4, 5). Mechanisms by which AID accesses S regions for CSR are well-understood (6, 7); but those by which AID accesses V(D)J exons for SHM are not (5).

In mature mouse B cells, transcription from an *Igh* V(D)J exon runs through C_H_-encoding Cμ, which encode the IgM heavy chains. Upon appropriate antigen or other activation, mature B cells undergo CSR, which replaces Cμ with one of 6 sets of C_H_s 100-200 kb downstream. Mouse C_H_s lie in the order 5′-V(D)J-Cμ-Cδ-Cγ3-Cγ1-Cγ2b-Cγ2a-Cε-Cα 3′ (3). Each C_H_ specifies an antibody class with different pathogen-elimination functions. Long repetitive switch (S) regions precede Cμ and downstream C_H_s (3). AID initiates CSR by deaminating cytosine residues within short sequence motifs, so-called RGYW nucleotide target motifs, the most robust being the palindromic AGCT sequence, that lie within donor Sμ and a downstream acceptor S region (8). The resulting AID-initiated DSBs are repaired through deletional end joining, which ligates the upstream Sμ break to a downstream acceptor S region break, thereby completing CSR (9). The 3′ *Igh* regulatory region (“3’RR”) is located at the downstream end of *Igh*, and contains 4 enhancers in a 30 kb region (10). The 3’RR is required for activating acceptor S region transcription, which targets AID, from flanking cytokine/activation-dependent promoters (3, 10–12). Cohesin-mediated loop extrusion plays a fundamental role in the CSR mechanism (1, 13). In resting B cells, the intronic enhancer (iEμ), the closely-linked Sμ and the downstream 30 kb long 3’RR function as dynamic, transcription-based loop anchors for cohesin-mediated loop extrusion between them (13). This process forms a nascent class-switch recombination-center (CSR-C) containing donor Sμ, iEμ and the 3’RR (13). Induced transcription, largely dependent on the 3’RR, from a targeted I-promoter that lies upstream of each acceptor S region renders that targeted region a dynamic extrusion impediment that leads to the targeted S region alignment with donor Sμ in the CSR-center (13). B cell activation also induces the expression of AID, which targets DSBs in aligned, transcribed S region donor and acceptor S regions, followed by their deletional end-joining that also is mediated by a cohesin-mediated extrusion-based mechanism within the CSR-C (1, 11, 13).

Mouse V(D)J exon SHM is targeted by transcription (3, 14). *Igh*, *Igκ* and *Igλ* light chain variable region exons contain three complementarity-determining region (CDR) sequences that encode antigen-contact sites that contain RGYW motifs, including AGCT motifs that, while at a much lower density than those in S regions, are frequent SHM targets (8). *Igh* and *Igκ* enhancers have also been implicated in SHM, potentially by promoting transcription through V(D)J exons to facilitate AID targeting (4, 15, 16).

Homozygous deletion of the 3’RR was found to abrogate low-level SHM of J_H_-iEμ intron sequences, but effects were not assayed in V(D)J exons (17–19). Homozygous 3’RR deletion reduced V(D)J transcription and also reduced BCR expression, which could indirectly affect the SHM process (17), leaving the precise role of the 3’RR in the SHM process to be fully determined. In mice, measurement of AID activity on a produtively rearranged *Igh* V(D)J allele, encoding VB1-8 heavy chain, versus a “non-productive” VB1-8 passenger allele that is identical except for an inactivating mutation, demonstrated that the V(D)J exon location is ‘‘privileged’’ for SHM targeting in GC B cells (8). Thus, these studies demonstrated that diverse sequences, including bacterial gene sequences, undergo SHM in place of the passenger VB1-8 sequence (8). Another long-standing question is how AID can be targeted at high levels to Sμ sequences but not in closely linked V(D)J exon sequences during CSR and, conversely, GC B cells undergoing SHM of V(D)J sequences do not undergo CSR (20). In this context, we proposed that in GC B cells, chromatin loop extrusion could directly juxtapose the 3’RR with the *Igh* variable region exon to establish a SHM center (SHM-C), that contributes to the privileged variable region exon location for SHM (13). Finally, a series of mouse studies have implicated SHM as predominantly occurring in the G1 cell cycle phase (21–26).

The human *IGH* C_H_ locus spans 300 kb downstream of the V(D)J exon and is organized differently from that of the mouse. Thus, the human C_H_ locus is organized into two tandemly duplicated clusters: 5′–V(D)J–Cμ–Cδ–Cγ3–Cγ1–Cα1–3’RR1–3’CBEs-1– Cγ2–Cγ4–Cα2–3’RR2–3’CBEs-2–3′ (10, 27) (Fig. 1A). Cμ and Cδ exons are only present in the upstream cluster, while each cluster contains Cγ, Cε, and Cα exons. Both clusters have a 3’RR and adjacent 3’CBEs, which could, in theory, separate them into distinct domains (10, 27). Similar to the mouse, multiple B cell–specific *cis*-regulatory elements are present in the distal human C_H_ locus, including iEμ, between J_H_s and Sμ, a putative element downstream of Cδ referred to as Eδ (27), and the V(D)J exon promoter known to be critical for SHM in mice (14). Cell line models played a fundamental role in efforts to discover and characterize the role of chromatin loop extrusion in V(D)J recombination and *Igh* CSR (1). In particular, mutations can be introduced and impacts characterized rapidly. A large body of work demonstrated the human Burkitt’s lymphoma-derived RAMOS GC B cell line to be a useful model for studies of SHM (27–32). In RAMOS cells, the *IGH* C_H_ locus exists in two distinct allelic configurations. One allele retains a productive V4-34/D3-10/J6 exon, which undergoes constitutive SHM in its endogenous chromosomal location; while the second allele is involved in a chromosomal translocation between Sμ and c-*MYC*, leading to dysregulated c-*MYC* expression and SHM (33). In this regard, we previously discovered that the 3’*RR* can activate c-*Myc* translocations several hundred kb upstream into Sμ in mouse B cell lymphomas (34). By analogy, RAMOS may activate SHM in *IGH* through a loop extrusion-based juxtaposition with the 3’RRs.

**Figure 1.**
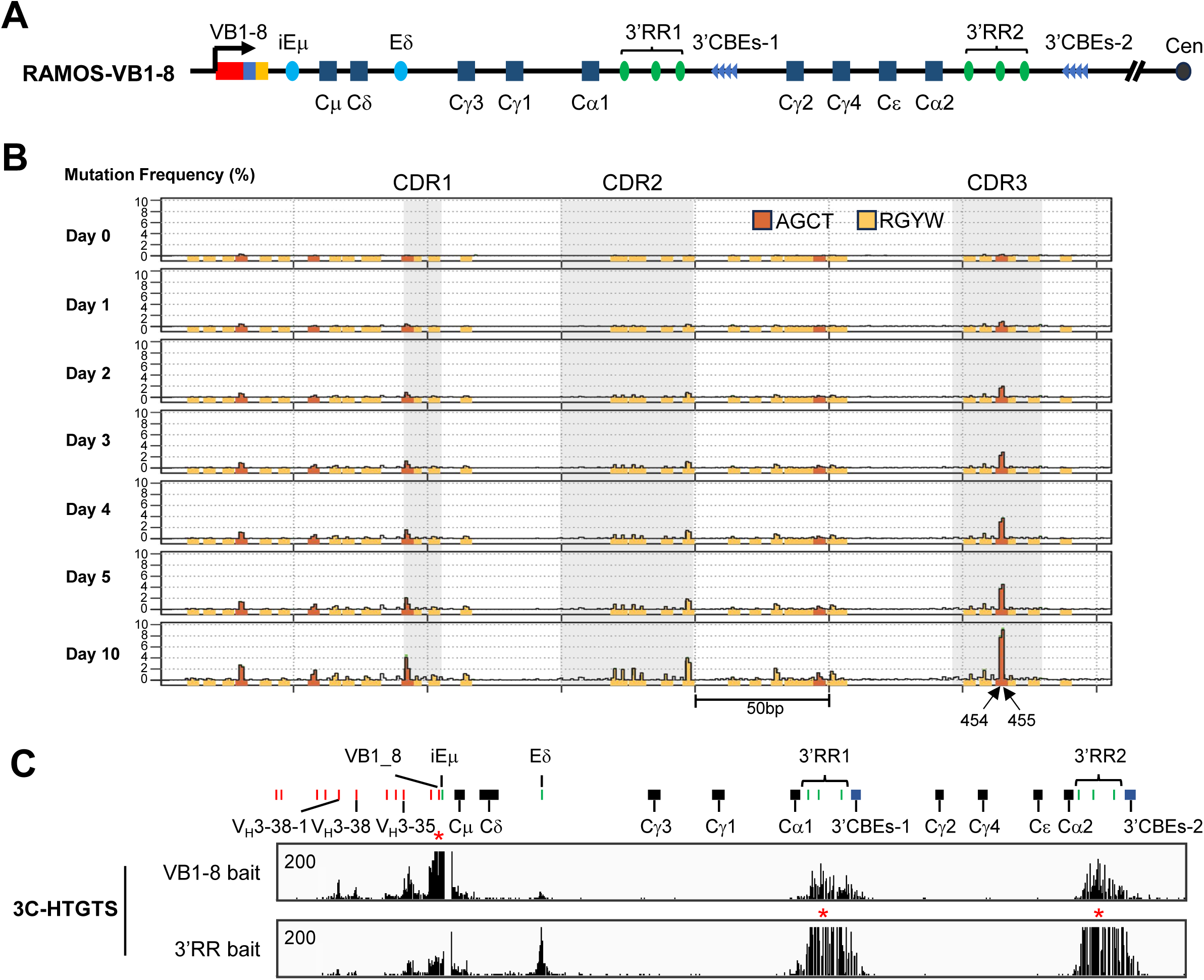
Chromatin organization and mutational landscape of the *IGH* locus in RAMOS-VB1-8 cells. (A) Schematic of the *IGH* constant region in the RAMOS-VB1-8 cell line. The endogenous rearranged human V(D)J was replaced with the mouse VB1-8 sequence. Individual enhancers within the 3′ regulatory regions (3′RRs) are depicted as green ovals. (B) Somatic hypermutation (SHM) profiles across the VB1-8 exon following doxycycline-induced AID expression at the indicated time points, as determined by SHM-Rep-Seq. The y-axis shows mutation frequency at each nucleotide position plotted as mean ± SEM from 3 independent repeats. Red and orange bars indicate AGCT and other RGYW motifs, respectively. (C) 3C-HTGTS interaction profiles across the 3′ *IGH* locus in RAMOS-VB1-8 cells, using VB1-8 and the 3′RRs as baits, respectively. Bait positions are indicated by red stars. All peaks are within the displayed scale range except for peaks at the bait sites. VB1-8 and upstream V_H_ segments are indicated by red lines; iEμ, Eδ, and individual 3′RR enhancers are indicated by green lines. Constant region exons are depicted as black boxes and 3′ CTCF-binding elements (3′CBEs) as blue boxes.

Decades ago, cycling RAMOS cells were found to constitutively undergo low-level SHM of their *IGH* and *IGL* variable region exons, and their translocated *c-MYC* gene (33), with SHM levels increased by introduction of a hyperactive AID mutant, AID7.3 (35). More recently, a further engineered RAMOS-based system was developed by deleting the endogenous AID gene (*AICDA)* and introducing a doxycycline-inducible AID7.3 cassette into the *AAVS1* locus, generating the AID7.3in cell line that was used to study the roles of enhancer-based transcription in stimulating SHM (31). The AID7.3in line was further engineered to generate the RASH line which contained a SHM reporter that was used to identify HMCES as a DNA repair factor involved in SHM (36). The RASH cell line and CH12/F3 mouse B cell lymphoma line were used to identify ELOF1 as a transcription elongation factor involved in SHM (37, 38). Another RAMOS system was developed in which endogenous *AICDA* was inactivated and the endogenous V(D)J region was replaced with other unmutated human V(D)J exons or the mouse VB1-8 V(D)J exon (32). In this system, SHM was induced by ectopic expression of a C-terminally truncated AID mutant, and SHM frequencies in the corresponding V exons were quantified by PCR-based sequencing and found to have patterns similar to VB1-8 in mouse GC B cells (32). However, in this study, roles of *cis*-regulatory element(s) in governing SHM were not addressed, leaving the relationship between SHM activity and transcriptional regulation an interesting unresolved question (32). Additional studies implicated the existence of an “*IGH* hub” structure involving interactions of the V(D)J exon with the 3’RRs, although potential roles of downstream interacting sequences in SHM including the 3’RRs were not reported (27). This hub structure was also detected 8 hours after RAD21 depletion in cycling RAMOS cells, suggesting the structure might be maintained in the absence of loop extrusion; but the potential for the maintained hub structure to support SHM after short-term RAD21 depletion was also not reported (27).

In the current work, we describe studies of an engineered AID7.3 RAMOS cell line in which we have inserted a VB1-8 allele in place of the endogenous RAMOS V(D)J exon. We also describe a further engineered version of this line that can be viably arrested for 10 days in the G1 phase of the cell cycle. Studies of these RAMOS lines have provided new insights into the existence of the putative SHM-C, the role of various elements including the 3’RRs in supporting V(D)J exon transcription versus SHM in the SHM-C, and the stability of the SHM-C complex and its ability to accumulate SHMs over multi-day periods in G1-arrested RAMOS cells in the presence or absence of RAD21.

## Results

### A modified human RAMOS cell line system to study SHM

To test mechanisms and elements that control SHM, we further modified the previously described human AID7.3in RAMOS-derived cell line, in which the endogenous *AICDA* gene was deleted and replaced with a doxycycline (Dox)-inducible hyperactive AID mutant (AID7.3) targeted to the AAVS1 locus (31). As the RAMOS *IGH* V4-34/D3-10/J6 expressed from its endogenous V4-34 promoter had already accumulated SHMs (28), we replaced it with the well-studied mouse VB1-8 exon that lacks any endogenous SHMs, allowing us to readily detect newly introduced SHMs. We previously characterized SHM patterns of the VB1-8 V(D)J exon in mouse GC B cells, as it served as the foundation for our mouse passenger-allele SHM studies (8). This unmutated, experimentally tractable V(D)J exon is transcribed from its own promoter and is also readily distinguishable from upstream human V_H_ segments in the various high-throughput assays used for our studies. For downstream studies described, we refer to the VB1-8 targeted AID7.3in cells with the endogenous *AICDA* gene deleted as the RAMOS-VB1-8 cells (Fig. 1A).

Following Dox-induced AID expression, we analyzed mutation profiles of the VB1-8 allele using our unbiased, high-throughput SHM-Rep-seq assay (39). Cycling RAMOS-VB1-8 cells accumulated substantial levels of SHM over a 10-day time course. During this time course, SHM frequencies steadily increased at AID hotspots including RGYW and AGCT motifs in both CDRs and framework regions (Fig. 1B). The overall SHM pattern was quite similar to that which we found for the mouse VB1-8 exon in mouse GC B cells (8). Overall, these studies established that RAMOS-VB1-8 cell lines accumulate SHMs at AID hotspots over the tested time course, and that levels and patterns resembled those of the same VB1-8 exons in normal mouse GC B cells that had accumulated 3-10 SHMs (8).

### Establishment of a SHM Center in RAMOS cells

To investigate the chromatin architecture associated with SHM targeting, we performed high-resolution 3C-HTGTS analyses on RAMOS-VB1-8 cell lines, baiting from the VB1-8 region and the HS1,2 region in the 3’RRs. Given the high resolution of 3C-HTGTS (40), we identified clear and robust interactions between the VB1-8 exon and both 3′RR1 and 3′RR2 (Fig. 1C, upper). These results revealed that the VB1-8 region forms broad interactions with both 3′RRs, and to a much lesser extent with both 3’CBEs. In addition, the VB1-8 region also interacted with a putative enhancer element termed Eδ (27), located approximately 20 kb downstream of Cδ, that is not conserved in the mouse C_H_ locus (10) (Fig. 1C, upper). The interactions of the two human 3’RRs with VB1-8 were distributed across HS3, HS1,2 and HS4 (Fig. 1C, upper). Reciprocally, baiting from the HS1,2 region in both 3’RRs (which are too homologous to distinguish sequence-wise), we confirmed that 3’RRs broadly interact with VB1-8 locale and those of iEμ and Eδ (Fig. 1C, lower). In contrast to mouse B cells activated for CSR, very few interactions were found with any other sequence regions including those containing S regions and C_H_ exons with VB1-8 and 3’RRs probes (Fig. 1C, upper and lower panels). Together, these findings establish a human SHM model system that recapitulates key features of *in vivo* SHM and reveal a network of long-range chromatin interactions linking the VB1-8 exon, iEμ, Eδ and the 3′RRs. These observations support the model that proposes these regulatory elements cooperatively organize a “SHM-C” that promotes efficient AID targeting and SHM, and that is structurally unique from that of a CSR-C that forms over the same region in B cells activated for CSR (13).

### Combined regulation of SHM by 3’RRs

To investigate the contribution of the two human 3′RRs to SHM, we generated a series of knockout cell lines, including lines with a 3′RR1-KO, a 3′RR2-KO, a 3′RR1/2- double KO (3’RR1&2-KO), and a larger 3′-region deletion encompassing both regulatory regions (“3’region-KO”) (Fig. 2A). Deletion of either individual 3′RR in the RAMOS-VB1-8 line caused at most a modest reduction in SHM frequency (Fig. 2B,S1).

**Figure 2.**
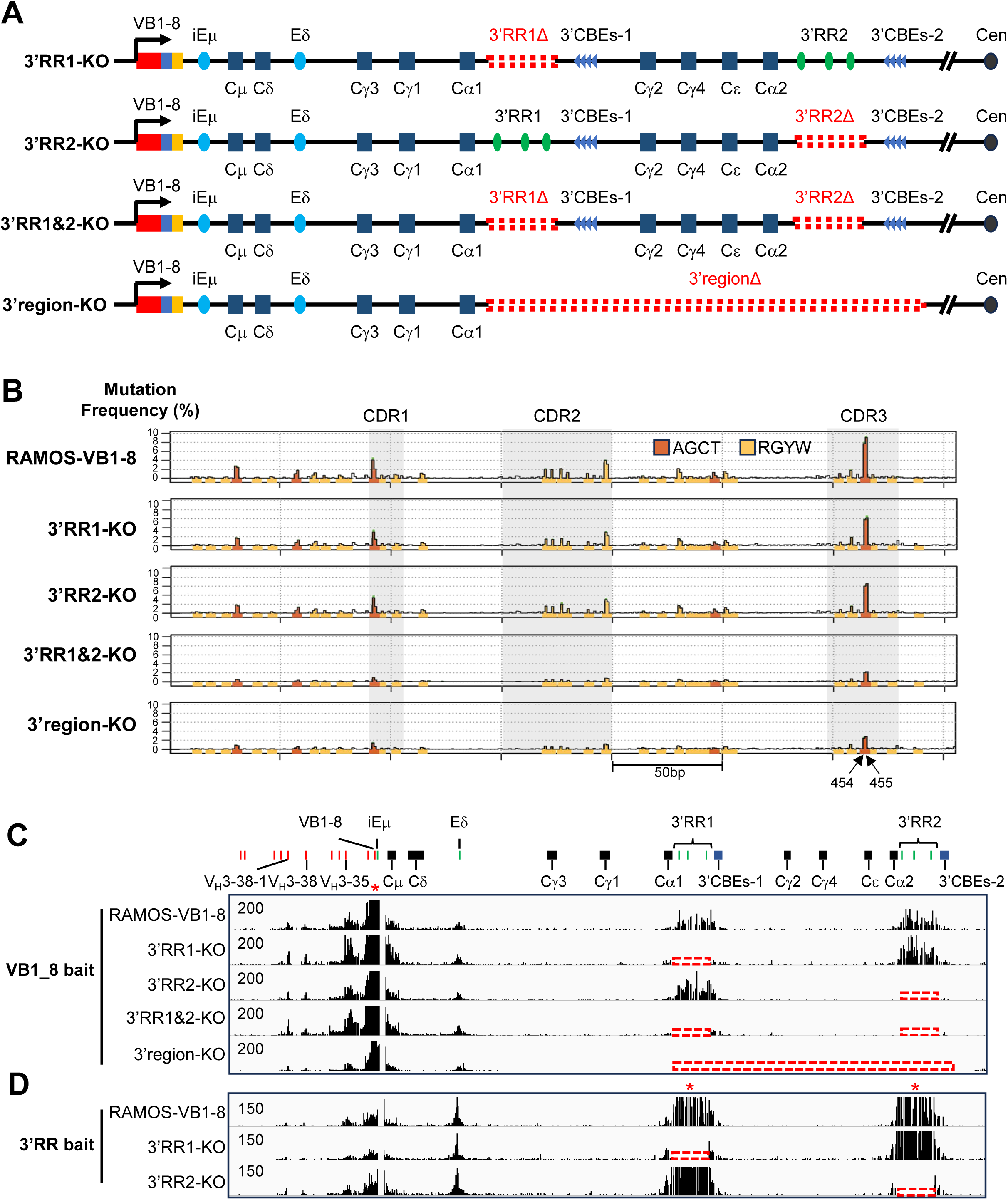
Impact of 3′RR deletions on SHM and chromatin interactions at the *IGH* locus. (A) Schematics of the *IGH* constant region in 3′RR1-KO, 3′RR2-KO, 3′RR1&2-KO, and 3′region-KO cell lines, illustrating the extent of each deletion. (B) SHM profiles across the VB1-8 exon in RAMOS-VB1-8, 3′RR1-KO, 3′RR2-KO, 3′RR1&2-KO, and 3′region-KO cells following 10 days of doxycycline-induced AID expression, as determined by SHM-Rep-Seq. Data is plotted as mean ± SEM from 3 independent repeats. RAMOS-VB1-8 and 3′RR1&2-KO SHM patterns are highly similar (two-sided Pearson’s *r* = 0.98, *P* = 3.9 x 10^-252^). RAMOS-VB1-8 and 3′region-KO SHM patterns are highly similar (two-sided Pearson’s *r* = 0.96, *P* = 3.04 x 10^-195^). (C) 3C-HTGTS interaction profiles across the 3′ *IGH* locus in RAMOS-VB1-8, 3′RR1-KO, 3′RR2-KO, 3′RR1&2-KO, and 3′region-KO cells, using VB1-8 as bait. (D) 3C-HTGTS interaction profiles across the 3′ *IGH* locus in RAMOS-VB1-8, 3′RR1-KO, and 3′RR2-KO cells, using the 3′RRs as bait.

Correspondingly, 3C-HTGTS-based chromatin interaction analyses revealed that the remaining intact 3′RR maintained robust long-range interactions with the VB1-8 region (Fig. 2C,D). Overall, these results demonstrated that each individual 3′RR is independently able to support efficient SHM in the absence of the other. In contrast, simultaneous deletion of both 3′RRs resulted in a substantial reduction in SHM frequency without affecting the pattern, but with still significant residual levels, confirming that the duplicated human 3′RRs function redundantly to promote efficient SHM (Fig. 2B, S1). Notably, deletion of the entire downstream region containing the two 3’RRs and both 3’CBEs did not further reduce residual SHM activity (Fig. 2B,S1). 3C-HTGTS analyses, baiting from the VB1-8 locus, reveal that VB1-8 interacts similarly with each 3’RR in the absence of the other and continues to interact with Eδ in the absence of both 3’RRs, suggesting a role of either iEμ, Eδ, or both in maintaining residual SHM activity (Fig. 2C).

### 3′RRs promote AID targeting independent of activating transcription

To test for potential mechanisms by which the 3’RRs promote SHM of V(D)J exons, we assessed their role in inducing or maintaining VB1-8 exon transcription, by performing PRO-seq in RAMOS-VB1-8 and the 3’region-KO cells in the absence of AID induction (Fig. 3A). In RAMOS-VB1-8 cells, transcription was initiated from the VB1-8 promoter and extended through the downstream intronic region containing iEμ, Sμ and the Cμ and Cδ exons (Fig. 3A, top panel). Further downstream, Eδ and both 3′RRs also exhibited robust transcriptional activity (27). Notably, these transcriptionally active regions corresponded closely to the chromatin interaction network identified by our 3C-HTGTS analyses, suggesting coordinated organization of transcriptional and structural regulatory elements within the SHM center (Fig. 1C). Surprisingly, the 3′region-KO cells retained robust transcription over the VB1-8, iEμ, Sμ and the Cμ and Cδ exons at levels comparable to those in RAMOS-VB1-8 cells (Fig. 3A). Within the VB1-8 locus, the rearranged V(D)J exon in both RAMOS-VB1-8 and 3′region-KO cells exhibited strong sense transcription accompanied by relatively low levels of antisense transcription (Fig. 3B,3C,S2A,S2B). Thus, deletion of the entire 3′ region had little effect on the quantitative transcriptional output across the VB1-8 region (Fig. 3D), despite causing a profound reduction in SHM frequency (Fig. 3B top, 3C top). These findings indicate that loss of the 3′RRs does not appreciably alter transcription levels prior to AID targeting and support findings that suggest the mouse 3′RRs can promote SHM through a transcription-level independent mechanism (17).

**Figure 3.**
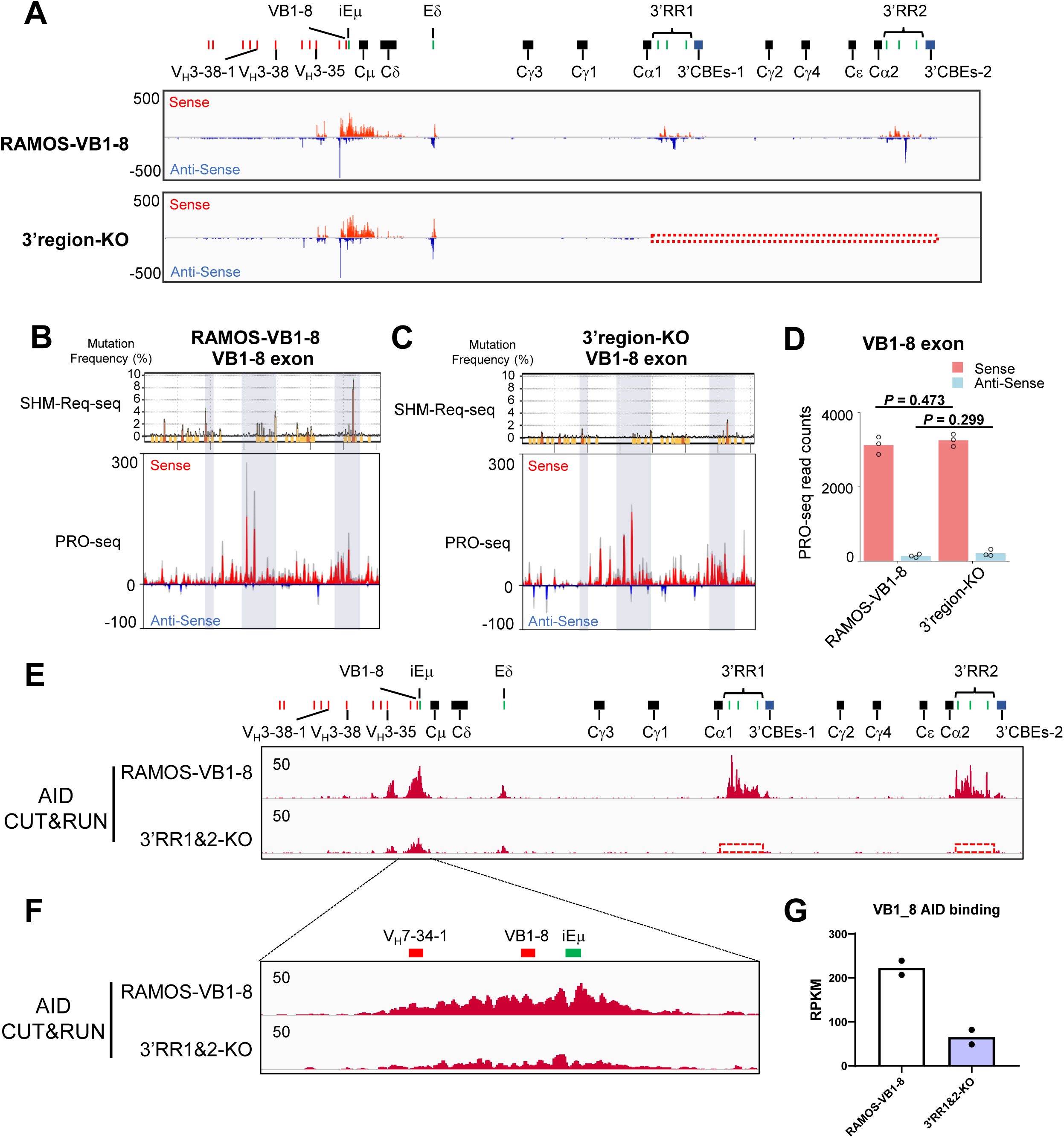
Transcriptional activity and AID occupancy at the *IGH* locus. (A) PRO-seq profiles across the *IGH* constant region in RAMOS-VB1-8 and 3′region-KO cells. (B–C) SHM profiles and normalized PRO-seq read counts across the VB1-8 exon in RAMOS-VB1-8 (B) and 3′ region-KO (C) cells. PRO-seq data represent the mean ± SD of three independent experiments. SHM profiles are plotted as mean ± SEM from 3 independent repeats. (D) normalized PRO-seq read counts across the VB1-8 exon in RAMOS-VB1-8 and 3′region-KO cells. Data represent the mean of three independent repeats. Statistical significance was assessed using a two-tailed unpaired Student’s t-test. (E–F) AID CUT&RUN profiles across the *IGH* constant region (E) and VB1-8 exon (F) in RAMOS-VB1-8 and 3′RR1&2-KO cells. (G) RPKM values of AID CUT&RUN signal across the VB1-8 exon in RAMOS-VB1-8 and 3′RR1&2-KO cells. Data represent the mean of two independent repeats.

To investigate whether the reduced SHM resulted from impaired AID recruitment, we performed AID CUT&RUN. AID accumulated robustly across the VB1-8 exon (Fig. 3E and F, top panels) as well as other transcribed sequences at the Eδ and 3’RRs in RAMOS *IGH* 3’ region (Fig. 3E and F, top panels). In contrast, deletion of both 3′RRs markedly reduced AID occupancy across the VB1-8 exon region (Fig. 3E and F, bottom panels, and Fig. 3G). Together, these results indicate that the 3′RRs facilitate efficient AID recruitment and/or retention at the rearranged V(D)J locus, beyond mechanisms involving transcription-level-mediated effects on increasing AID substrate targeting ability (41).

### Robust AID-induced SHM over a 10-day time-course in G1-arrested RAMOS Cells

Various studies have indicated that AID initiates SHM during the G1 cell cycle phase in GC B cells (22–25). We sought to test the ability of AID to induce SHMs in G1-arrested RAMOS cells. We found that the CDK4/6 inhibitor PD0332991 efficiently arrested RAMOS cells in the G1 phase, thereby providing a system to examine SHM under G1-arrest conditions (Fig. 4A). However, prolonged G1 arrest resulted in a substantial loss of cell viability over the course of PD0332991 treatment (Fig. 4B). To improve viability during extended G1 arrest, we introduced a *Bcl2* overexpression cassette into the AAVS1 locus of the RAMOS-VB1-8 cells (Fig. 4C,D). Hereafter, we refer to the Bcl2-overexpressing RAMOS-VB1-8 cells as RAMOS-VB1-8-Bcl2 cells.

**Figure 4.**
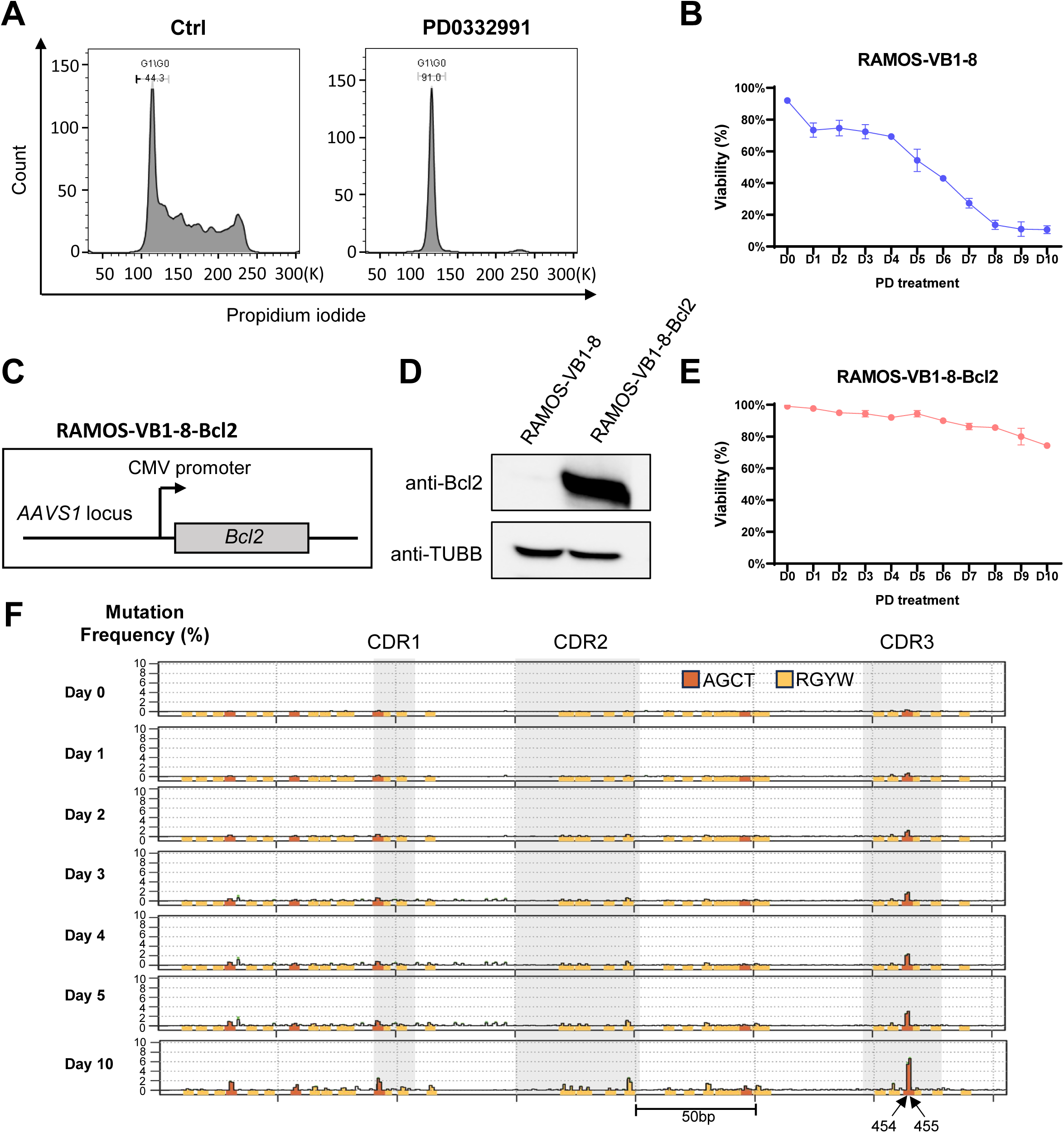
Establishment and validation of a cell cycle-arrested system for SHM analysis. (A) Flow cytometry of propidium iodide-stained RAMOS-VB1-8 cells with or without PD0332991 treatment. (B) Cell viability of RAMOS-VB1-8 cells following 10 days of PD0332991 treatment. Data represent the mean ± SD of three independent repeats. (C) Schematic of the AAVS1 locus depicting the integrated CMV-*Bcl2* overexpression cassette in RAMOS-VB1-8-Bcl2 cells. (D) Western blot analysis of Bcl2 protein levels in RAMOS-VB1-8 and RAMOS-VB1-8-Bcl2 cells. Raw data is shown in Figure S8A. (E) Cell viability of RAMOS-VB1-8-Bcl2 cells following 10 days of PD0332991 treatment. Data represent the mean ± SD of three independent repeats. (F) SHM profiles across the VB1-8 exon in G1-arrested RAMOS-VB1-8-Bcl2 cells following doxycycline-induced AID expression at the indicated time points, as determined by SHM-Rep-Seq. Data is plotted as mean ± SEM from three independent repeats.

RAMOS-VB1-8-Bcl2 cells maintained high viability even after 10 days of PD0332991 treatment (Fig. 4E). Strikingly, G1-arrested RAMOS-VB1-8-Bcl2 cells remained capable of undergoing robust SHM following AID induction. Moreover, the G1-arrested RAMOS-VB1-8-Bcl2 cells accumulated SHMs over the 10-day test period with a similar pattern to that of cycling RAMOS-VB1-8 cells (Fig. 1B and 4F). The final SHM frequencies were only modestly reduced compared with those in cycling cells (Fig. 1B, 4F, and S3).

### Robust SHM in 10-day time-course in absence of RAD21 in G1-arrested RAMOS Cells

To begin to test the proposed role of cohesin-mediated loop extrusion in assembly of the SHM-C (13), we sought to deplete the RAD21 cohesin ring component in RAMOS cells. RAD21 inactivation generates a rapid decrease in viability in cycling mammalian cells (42, 43). In that regard, we previously reported that we can viably deplete numerous different cohesin components, including RAD21, in G1-arrested, v-*Abl*-transformed pro-B cells that ectopically express Bcl2 (44, 45). Based on these findings, we generated a homozygous RAD21 degron cell line in RAMOS-VB1-8-Bcl2 cells by tagging both endogenous RAD21 alleles with a FKBP12^F36V^ degron cassette (Fig. S4A) (46). Hereafter, we refer to the RAMOS-VB1-8-Bcl2 cells with homozygous RAD21 degron as RAD21-degron cells. Western blot analysis confirmed efficient depletion of RAD21 protein in the RAD21-degron cells within 6 h of dTAGV-1 treatment (Fig. S4B). Under G1 arrest, RAD21-depleted cells maintained nearly 80% viability for 4 days and approximately 60% viability at 10 days (Fig. S4C), providing a suitable system for investigating the long-term effect of RAD21 depletion on SHM and the SHM-C architecture.

To define the genomic distribution of cohesin, we performed CUT&RUN for RAD21 (Fig. 5A). Before dTAGV-1 treatment, RAD21 was highly enriched at both 3′CBEs, consistent with their role as loop anchors. Lower levels of RAD21 accumulation were detected at the 3′RRs, Eδ, and several sites within the upstream V region, suggesting that cohesin contributes to the formation of the SHM center. As expected, chromatin-associated RAD21 accumulation was essentially absent following 6 h of dTAGV-1 treatment (Fig. 5A). To examine effects of RAD21 depletion on chromatin interactions genome-wide, RAD21-degron cells were treated with or without dTAGV-1 for 6 h before G1 arrest and AID induction, and after 4 days of RAD21 depletion, cells were harvested for Hi-C analysis. The *IGH* 3′ region contains two sets of homologous constant regions with high sequence similarity, and contains repetitive sequences in the switch regions, 3′RRs, and 3’CBEs, which creates challenges for mapping Hi-C reads in this region (Fig. S4D). Even so, these studies showed that chromatin loops and topologically associating domains (TADs) were essentially eliminated genome-wide (Fig. S4E), confirming the effective depletion of RAD21.

**Figure 5.**
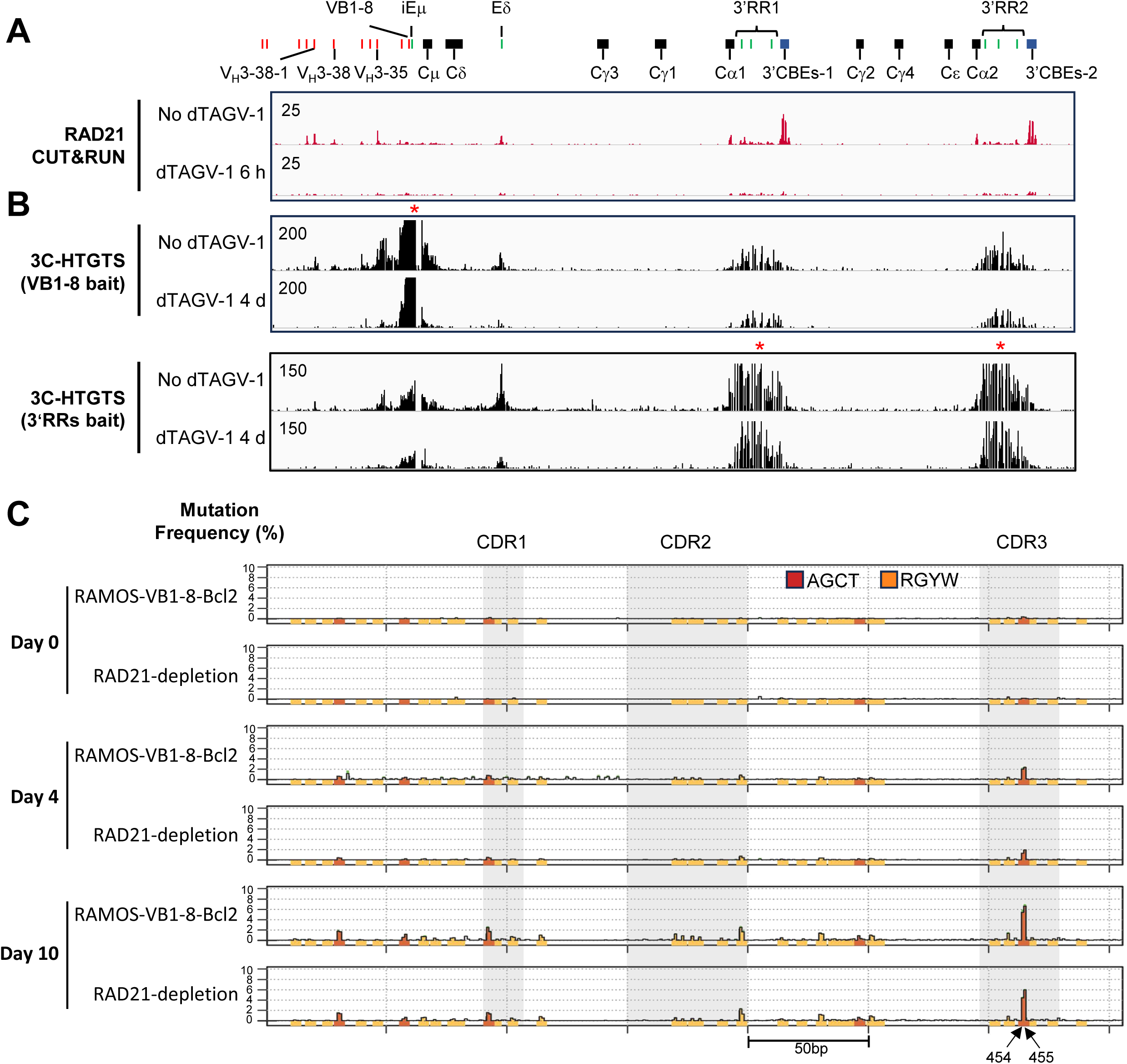
Effects of RAD21 depletion on chromatin organization and SHM at the *IGH* locus. (A) CUT&RUN profiles of RAD21 occupancy across the *IGH* locus in RAD21-degron cells treated with or without dTAGV-1 for 6 hours. (B) 3C-HTGTS interaction profiles across the 3′ *IGH* locus in RAD21-degron cells treated with or without dTAGV-1 for 4 days, using VB1-8 and 3′RRs as baits, respectively. (C) SHM profiles across the VB1-8 exon in G1-arrested RAMOS-VB1-8-Bcl2 and RAD21-depleted cells following doxycycline-induced AID expression at the indicated time points, as determined by SHM-Rep-Seq. Data is plotted as mean ± SEM from three independent repeats. RAMOS-VB1-8-Bcl2 and RAD21-depletion SHM patterns are highly similar (Day 4: two-sided Pearson’s *r* = 0.86, *P* = 4.59 x 10^-106^; Day 10: two-sided Pearson’s *r* = 0.99, *P* = 2.44 x 10^-292^).

To determine the contribution of cohesin to SHM-associated chromatin architecture, we next examined chromatin interactions within the *IGH* locus at high resolution using 3C-HTGTS with the VB1-8 region and 3′RRs as bait four days after RAD21 depletion. Remarkably, despite the absence of most chromatin loop domains genome-wide, long-range interactions between VB1-8 and the downstream 3′RRs were still substantially preserved after four days of RAD21 depletion (Fig. 5B, S4F). This finding indicates that the VB1-8–3′RR-based SHM-C is largely maintained intact for at least four days in the absence of cohesin-mediated loop extrusion in G1-arrested RAMOS cells (Fig. 5B, S4F). In contrast, interactions between VB1-8 and immediately upstream regions were substantially decreased (Fig. 5B, S4F). To determine whether cohesin loss affected transcription, we performed PRO-seq following four days of RAD21 depletion. Transcription across the IGH C_H_ locus was largely unchanged (Fig. S5A), and transcription level from the VB1-8 locus was only slightly increased (Fig. S5B-D). Finally, consistent with the persistence of both VB1-8–3′RR-based SHM-C structure and its transcriptional activity, SHM-rep-seq analysis showed that RAD21 depletion in the G1-arrested RAD21-degron cells had essentially no effect on SHM accumulation in the VB1-8 exon, as compared to that of the G1-arrested RAMOS-VB1-8-Bcl2 cells or RAD21-degron cells without dTAGV-1 treatment. (Fig. 5C, S6A, S6B). Together, these results demonstrate that ongoing cohesin-mediated loop extrusion is largely dispensable for maintenance of the architectural organization, transcriptional activity, and functional SHM activity of the SHM-C over a multi-day period in G1-arrested RAMOS cells.

## Discussion

The human RAMOS germinal center B cell lymphoma line has for many decades provided an important *in vitro* model system for studying the mechanism of SHM (27–32). For our current studies, we made several modifications of this line. We further modified the AID7.3in RAMOS cell line, in which the endogenous AID gene had been deleted and an inducible hyperactive AID7.3 mutant had been introduced (31). The first modification we made was to replace the endogenous *IGH* V(D)J exon with an unmutated mouse VB1-8 V_H_(D)J_H_ exon (Fig. 1A), a V(D)J exon for which we previously characterized the SHM pattern in depth in response to immunization in mouse GC B cells (8). We named the resulting cell line the RAMOS-VB1-8 line. We used our high-throughput SHM-Rep-Seq method to characterize the SHM level and pattern upon AID induction over a 10-day time course in the RAMOS-VB1-8 line (Fig. 1B). In this line, the VB1-8 exon accumulated SHMs mainly focused on AID-targeting motifs, most robustly the palindromic AGCT canonical AID target motif, with a pattern similar to that which we reported for GC B cells (Fig. 1B) (8). The SHMs accumulated over the 10-day time course reaching moderate levels that were similar to those of mouse GC B cells that accumulated 3-10 SHMs per sequence in pseudo kinetic assays (Fig.1B) (8). Mouse GC SHM levels reach much higher levels than those we observed in the RAMOS-VB1-8 cell line studies (8). While we are unable at this time to explain the lower levels of SHMs accumulated over our 10-day RAMOS time course, there are several possibilities. One is that AID activity may be rate-limiting in RAMOS cells for several potential reasons discussed below. Another is that high-level accumulation of SHMs in mouse GC B cells can involve multiple rounds of B cell re-entry into the GC over the course of immunization (47). In theory, we could address this question by attempting several rounds of AID induction, particularly with the G1-arrestable model that we have generated.

We employed our robust and high-resolution 3C-HTGTS assay to test our hypothesis that an SHM-C, analogous to the CSR-C, provides a privileged location for the accumulation of SHMs (Fig. 1C) (13). Indeed, similar to what we found for the CSR-C in mouse activated B cells, we found that the V(D)J exon in RAMOS cells robustly interacts across the 20-kb length of each 3’RR1 and 3’RR2, and also with other putative elements, such as the putative Eδ enhancer (Fig. 1C) (13, 27). To test the potential roles of the two human 3’RR sequences in supporting SHM in the human SHM-C, we deleted 3’RR1, 3’RR2, or both in the RAMOS-VB1-8 line (Fig. 2A). Deletion of either of the 3’RRs had no obvious effect on the interactions of VB1-8 with the other remaining 3’RR and had, at most, modest effects on SHM over the 10-day time course (Fig. 2B,C). On the other hand, deletion of both 3’RRs or the entire region containing them and their associated 3’CBEs reduced SHM levels by approximately 4-fold but did not alter the SHM pattern (Fig. 2B,S1). These overall findings indicated that either human 3’RR can form a functional SHM-C in the absence of the other, but in the absence of both, SHM is greatly reduced.

PRO-seq analysis showed that complete deletion of the 3′RRs and associated sequences did not markedly alter the transcription levels across the VB1-8 exon, despite the major effect that their deletion had on SHM (Fig. 3A-D). We note that this observation on transcription levels does not exclude changes in other transcriptional features in the RAMOS-VB1-8 versus the 3′RR region KO cell line, such as RNA polymerase II (Pol II) pausing or stalling within the VB1-8 exon. In this regard, previous studies have suggested that Pol II pausing/stalling facilitates AID targeting (15, 37, 38, 48, 49). Consistent with this idea, our CUT&RUN analysis showed that the deletion of both 3′RRs reduced AID occupancy across the VB1-8 exon, supporting a role for the 3′RRs in promoting AID recruitment (Fig. 3E-G) (15). These observations are also consistent with our SHM-C model, in which chromatin loop extrusion brings the 3′RRs into proximity with the V(D)J region, thereby facilitating the recruitment or retention of AID and/or AID-associated transcriptional regulators (13, 15, 37, 38, 50, 51). Together, these findings raise the possibility that, although overall transcriptional output remains unchanged, the dual 3′RR deletion alters transcriptional dynamics (e.g., Pol II pausing/stalling) and/or the recruitment of transcription-associated factors, which could impair both AID accumulation and its transcription-targeted activity across the *IGH* V(D)J exon. Recent studies demonstrated that S region transcripts recruit AID to the *Igh* CSR-C to initiate CSR (7). As V(D)J exons lack dense arrays of transcribed AID targeting motifs present in S regions, any role for V(D)J exon transcripts in recruiting AID into a SHM-C would likely employ a different, yet unknown, mechanism.

The 3′CBEs have been proposed to serve as a major loop anchor within the mouse *Igh* locus (13, 52). We note that the RAMOS VB1-8 exon retained robust interactions with the 3′RRs even though the 3′CBE-1 cluster is positioned between the two 3′RRs (Fig. 1C). This finding suggests that the intervening 3′CBE-1 cluster does not block communication between the VB1-8 exon l and the 3′RRs. One possibility is that these CBEs anchor convergent chromatin loops, thereby bringing the VB1-8 exon into close spatial proximity with both 3′RRs (See Model in Fig. S7). Another possibility is that the impediment activity of the human 3’CBEs, which only have 4 CBE copies, is less than that of the mouse, which have 10 CBE copies (1). It is also theoretically possible that the 3′CBE impediment activity is functionally attenuated or neutralized in RAMOS cells, as previous studies have shown that CBE function can be modulated during B-cell development (45).

A major innovation of our RAMOS-VB1-8 model was our finding that enforced Bcl2 overexpression allowed RAMOS-VB1-8-Bcl2 cells to be viably G1-arrested for up to 10 days in the G0/G1 stage of the cell cycle (Fig. 4A-E). When these cells were G1-arrested for 10 days after AID induction, they accumulated VB1-8 mutations to a similar level and pattern as cycling RAMOS cells over the 10-day time course (Fig. 4F). This finding is of interest given prior cell cycle studies of AID activity in B cells. In this regard, it has been suggested that nuclear AID is more stable during the G1 stage (22). In addition, AID has been shown to undergo active nuclear import (53). Recent *in vivo* studies have reported that GC B cells exhibit higher mutation efficiency during the G0/G1 stage (24, 25). Yet, whether AID is actually enriched in the nucleus during G1 arrest remains controversial (23, 53, 54). We note that our G1-arrested RAMOS-VB1-8-Bcl2 cells did not display higher SHM efficiency than cycling cells (Fig. S3). Potential differences between our RAMOS findings and those of other studies in mouse GC B cells may reflect differences in cell types, experimental systems, or imaging strategies.

However, our G1-arrestable RAMOS system should provide a useful system for pursuing single molecule/single cell imaging studies (7) to address relationships between nuclear accumulation of AID in the SHM-C and influences on SHM activity across the cell cycle.

Finally, we have shown that our RAMOS-VB1-8-Bcl2 cell SHM model, similar to our earlier findings with the G1-arrested *v-Abl* cell model (44, 45), allows us to viably deplete RAD21 in G1-arrested cells to begin to test the potential roles of cohesin complex factors in the SHM process (Fig. S4). Analogies to the CSR-C support the notion that the SHM-C may be formed through cohesin-mediated loop extrusion between the 3′RRs and upstream transcription complexes in the general vicinity of the V(D)J exon (13). In this regard, loop extrusion also has been postulated to be implicated in bringing enhancers and AID targets together in other chromosome loop domains (30). We note that four days of cohesin depletion did not abolish the VB1-8/3′RR interaction, suggesting that maintenance of SHM-C interactions in G1-arrested cells is largely independent of ongoing cohesin-mediated loop extrusion (Fig. 5). A similar persistence of the VB1-8–3′RR interaction was previously noted following 8 hours of RAD21 depletion in cycling cells (27). Additional factors might therefore contribute to the establishment and/or maintenance of the three-dimensional architecture of the *Igh* locus, which will be the focus of ongoing studies (30, 55, 56).

## Materials and Methods

### Cell culture

RAMOS B cells were cultured in RPMI-1640 medium, supplemented with 10% FBS and 1X Penicillin-Streptomycin-Glutamine. For induction of AID expression, cells were treated with 200 ng/ml doxycycline (Dox) for the indicated durations. For PD0332991-induced G1 arrest, cells were treated with 10 μM PD0332991 for the indicated durations. For RAD21 depletion, RAD21-degron cells were first treated with 500 nM dTAGV-1 for 6 hours, followed by treatment with 10 μM PD0332991 and 200 ng/ml Dox for the indicated durations prior to collection and analysis.

### Generation of mutant RAMOS cell lines

All cell lines were generated using a CRISPR/Cas9 strategy. For cell lines targeting the IGH constant region specifically, 3C-HTGTS using the c-*MYC* gene as bait was performed to confirm that editing occurred exclusively at the VB1-8 allele rather than the translocated allele. sgRNA sequences used for cell line generation are listed in Supplementary Table 1.

### Flow cytometric analysis

The distribution of cells across the cell cycle was assessed by propidium iodide (PI) staining, followed by flow cytometry. Cells were treated with or without 10 μM PD0332991 for 24 hours, then collected and washed once with 1x DPBS. Cells were fixed in 70% ethanol at −20°C for 24 hours, pelleted by centrifugation, and resuspended in 1x DPBS containing 100 μg/ml RNase A. Following incubation at 37°C for 30 minutes, cells were pelleted, resuspended in 1x DPBS containing 10 μg/ml PI, and incubated on ice for 10 minutes. Samples were then analyzed on a BD FACS Calibur flow cytometer (BD Biosciences). Data were analyzed using FlowJo software (v10). Intact cells were gated based on FSC-A/SSC-A parameters, and single cells were isolated using an FSC-A/FSC-H gate. Cell cycle distribution was determined based on PI fluorescence intensity.

### SHM-Rep-Seq and data analysis

SHM-Rep-Seq libraries were prepared using VB1-8 locus as bait, and the procedure was conducted as previously described (57).

Libraries were sequenced on a NextSeq 2000 (Illumina) with 300-bp paired-end reads. SHM-rep-seq data was analyzed as previous described(8). Bio primers and red primers are listed in Supplementary Table 2.

### 3C-HTGTS and data analysis

3C-HTGTS libraries were prepared as described(40). Libraries were sequenced on a NextSeq 2000 (Illumina) with 300-bp paired-end reads. Bio primers and red primers are listed in Supplementary Table 2. The 3C-HTGTS data was analyzed as previously described by aligning to hg38 reference genome containing the engineered *IGH* allele (40). For each figure, libraries were normalized to the same number of junctions prior to visualization.

### PRO-Seq and data analysis

PRO-seq experiments were performed as previously described (58). Libraries were sequenced on a NextSeq 2000 (Illumina) with 50-bp paired-end reads. PRO-seq data were aligned to a modified hg38 reference genome containing the engineered IGH allele using Bowtie2 (v2.4.2). BAM files were sorted and indexed using SAMtools (v0.1.19). Strand-specific normalized PRO-seq read counts were generated from the processed BAM files using BEDTools (v2.29.0) and bamCoverage (v3.3.1) with a 1-bp bin size. The normalized PRO-seq read counts were visualized using Integrative Genomics Viewer (2.19.2) and analyzed using Python (3.7.4) scripts.

### CUT&RUN and data analysis

CUT&RUN experiment was performed using a CUT&RUN Library Prep Kit (EpiCypher, 14-1048) according to the manufacturer’s instructions. Libraries were sequenced on a NextSeq 2000 (Illumina) with 150-bp paired-end reads. The antibodies used for CUT&RUN are listed in Supplementary Table 3.

CUT&RUN data were aligned to a modified hg38 reference genome containing the engineered IGH allele using Bowtie2 (v2.4.2). BAM files were sorted and indexed using SAMtools (v0.1.19). Normalized coverage tracks were generated from the resulting BAM files using bamCoverage (v3.3.1) and CPM normalization.

### Hi-C and data analysis

Hi-C experiments were performed using an Arima-HiC Kit (Arima, A510008) according to the manufacturer’s instructions. Sequencing libraries were generated with the NEBNext Ultra II DNA Library Prep Kit (NEB, E7645) according to the manufacturer’s guidelines. Libraries were sequenced on a NextSeq 2000 (Illumina) with 150-bp paired-end reads. Hi-C sequencing data were processed using the CPU version of Juicer v1.6 (https://github.com/aidenlab/juicer). Reads were aligned to the modified hg38 reference genome containing the engineered IGH allele using the default Juicer alignment workflow. Valid interaction pairs with mapping quality (MAPQ) ≥30 were retained for downstream analyses, and contact maps were visualized in Juicebox.

### Statistical analysis

Statistical analyses in Figs. 3D, S1, S3, S4F, S5D and S6A were performed using a two-tailed unpaired Student’s *t*-test. Statistical analyses in Figs. 2B and 5C were performed using a two-sided Pearson correlation test.

## Acknowledgments

We thank the F.W.A. laboratory members for comments and discussions about this study. We thank Jayanta Chaudhuri for sharing the AID CUT&RUN antibody. This work was supported by NIH grant CSR/SHM 5R37AI077595. F.W.A. is an investigator of the Howard Hughes Medical Institute.

## Data Availability

All the sequencing data will be deposited at public dataset after accept.

## References

1. Y. Zhang, X. Zhang, H. Q. Dai, H. Hu, F. W. Alt, The role of chromatin loop extrusion in antibody diversification. Nat Rev Immunol 22, 550–566 (2022).

2. M. Muramatsu et al., Class Switch Recombination and Hypermutation Require Activation-Induced Cytidine Deaminase (AID), a Potential RNA Editing Enzyme. Cell 102, 553–563 (2000).

3. W. T. Yewdell, K. C. Fernandez, S. M. Downs-Canner, J. Chaudhuri, The Molecular Logic of Immunoglobulin Heavy Chain Class Switch Recombination. Annu Rev Immunol 44, 527–551 (2026).

4. R. Casellas et al., Mutations, kataegis and translocations in B cells: understanding AID promiscuous activity. Nat Rev Immunol 16, 164–176 (2016).

5. J. Ridani, P. Barbulescu, A. Martin, J. M. D. Noia, Somatic Hypermutation. T. Honjo, M. Reth, A. Radbruch, F. Alt, A. Martin, Eds., Molecular Biology of B cells (Elsevier, 2023), pp. 235–256.

6. Y. Zhang et al., The fundamental role of chromatin loop extrusion in physiological V(D)J recombination. Nature 573, 600–604 (2019).

7. M. Mikhova et al., A dynamic RNA hub facilitates activation-induced cytidine deaminase recruitment to the immunoglobulin heavy-chain locus. Mol Cell 10.1016/j.molcel.2026.06.012 (2026).

8. L. S. Yeap et al., Sequence-Intrinsic Mechanisms that Target AID Mutational Outcomes on Antibody Genes. Cell 163, 1124–1137 (2015).

9. J. Dong et al., Orientation-specific joining of AID-initiated DNA breaks promotes antibody class switching. Nature 525, 134–139 (2015).

10. E. Pinaud et al., The IgH locus 3’ regulatory region: pulling the strings from behind. Adv Immunol 110, 27–70 (2011).

11. J. Chaudhuri et al., Transcription-targeted DNA deamination by the AID antibody diversification enzyme. Nature 422, 726–730 (2003).

12. E. Pinaud et al., Localization of the 3’ IgH Locus Elements that Effect Long Distance Regulation of Class Switch Recombination. Immunity 15, 187–199 (2001).

13. X. Zhang et al., Fundamental roles of chromatin loop extrusion in antibody class switching. Nature 575, 385–389 (2019).

14. Y. Fukita, H. Jacobs, K. Rajewsky, Somatic Hypermutation in the Heavy Chain Locus Correlates with Transcription. Immunity 9, 105–114 (1998).

15. A. Tarsalainen et al., Ig Enhancers Increase RNA Polymerase II Stalling at Somatic Hypermutation Target Sequences. J Immunol 208, 143–154 (2022).

16. J. M. Buerstedde, J. Alinikula, H. Arakawa, J. J. McDonald, D. G. Schatz, Targeting of somatic hypermutation by immunoglobulin enhancer and enhancer-like sequences. PLoS Biol 12, e1001831 (2014).

17. P. Rouaud et al., The IgH 3’ regulatory region controls somatic hypermutation in germinal center B cells. J Exp Med 210, 1501–1507 (2013).

18. A. Saintamand et al., Deciphering the importance of the palindromic architecture of the immunoglobulin heavy-chain 3’ regulatory region. Nat Commun 7, 10730 (2016).

19. C. L. Morvan, E. Pinaud, C. Decourt, A. Cuvillier, M. Cogne, The immunoglobulin heavy-chain locus hs3b and hs4 3’ enhancers are dispensable for VDJ assembly and somatic hypermutation. Blood 102, 1421–1427 (2003).

20. J. A. Roco et al., Class-Switch Recombination Occurs Infrequently in Germinal Centers. Immunity 51, 337–350 e337 (2019).

21. G. Sharbeen, C. W. Yee, A. L. Smith, C. J. Jolly, Ectopic restriction of DNA repair reveals that UNG2 excises AID-induced uracils predominantly or exclusively during G1 phase. J Exp Med 209, 965–974 (2012).

22. Q. Le, N. Maizels, Cell Cycle Regulates Nuclear Stability of AID and Determines the Cellular Response to AID. PLoS Genet 11, e1005411 (2015).

23. Q. Wang et al., The cell cycle restricts activation-induced cytidine deaminase activity to early G1. J Exp Med 214, 49–58 (2017).

24. J. Pae et al., Transient silencing of hypermutation preserves B cell affinity during clonal bursting. Nature 641, 486–494 (2025).

25. J. Merkenschlager et al., Regulated somatic hypermutation enhances antibody affinity maturation. Nature 641, 495–502 (2025).

26. A. Faili et al., AID-dependent somatic hypermutation occurs as a DNA single-strand event in the BL2 cell line. Nat Immunol 3, 815–821 (2002).

27. U. E. Schoeberl et al., Regulation of somatic hypermutation by higher-order chromatin structure. Mol Cell 10.1016/j.molcel.2025.06.003 (2025).

28. J. E. Sale, M. S. Neuberger, TdT-Accessible Breaks Are Scattered over the Immunoglobulin V Domain in a Constitutively Hypermutating B Cell Line. Immunity 9, 859–869 (1998).

29. F. N. Papavasiliou, D. G. Schatz, Cell-cycle-regulated DNA doublestrand breaks in somatic hypermutation of immunoglobulin genes. Nature 408, 216–220 (2000).

30. F. Senigl et al., Topologically Associated Domains Delineate Susceptibility to Somatic Hypermutation. Cell Rep 29, 3902–3915 e3908 (2019).

31. R. K. Dinesh et al., Transcription factor binding at Ig enhancers is linked to somatic hypermutation targeting. Eur J Immunol 50, 380–395 (2020).

32. U. E. Schoeberl et al., Somatic hypermutation patterns in immunoglobulin variable regions are established independently of the local transcriptional landscape. eLife 14:RP106566 (2025).

33. M. Bemark, M. S. Neuberger, The c-MYC allele that is translocated into the IgH locus undergoes constitutive hypermutation in a Burkitt’s lymphoma line. Oncogene 19, 3404–3410 (2000).

34. M. Gostissa et al., Long-range oncogenic activation of Igh-c-myc translocations by the Igh 3’ regulatory region. Nature 462, 803–807 (2009).

35. M. Wang, Z. Yang, C. Rada, M. S. Neuberger, AID upmutants isolated using a high-throughput screen highlight the immunity/cancer balance limiting DNA deaminase activity. Nat Struct Mol Biol 16, 769–776 (2009).

36. L. Wu et al., HMCES protects immunoglobulin genes specifically from deletions during somatic hypermutation. Genes Dev 36, 433–450 (2022).

37. L. Wu et al., Transcription elongation factor ELOF1 is required for efficient somatic hypermutation and class switch recombination. Mol Cell 10.1016/j.molcel.2025.02.007 (2025).

38. P. Dai et al., Transcription-coupled AID deamination damage depends on ELOF1-associated RNA polymerase II. Mol Cell 10.1016/j.molcel.2025.02.006 (2025).

39. H. Chen et al., BCR selection and affinity maturation in Peyer’s patch germinal centres. Nature 582, 421–425 (2020).

40. S. Jain, Z. Ba, Y. Zhang, H. Q. Dai, F. W. Alt, CTCF-Binding Elements Mediate Accessibility of RAG Substrates During Chromatin Scanning. Cell 174, 102–116 e114 (2018).

41. B. Laffleur et al., AID-induced remodeling of immunoglobulin genes and B cell fate. Oncotarget 5, 1118–1131 (2013).

42. S. S. P. Rao et al., Cohesin Loss Eliminates All Loop Domains. Cell 171, 305–320 e324 (2017).

43. M. Gabriele et al., Dynamics of CTCF- and cohesin-mediated chromatin looping revealed by live-cell imaging. Science 376, 496–501 (2022).

44. Z. Ba et al., CTCF orchestrates long-range cohesin-driven V(D)J recombinational scanning. Nature 586, 305–310 (2020).

45. H. Q. Dai et al., Loop extrusion mediates physiological Igh locus contraction for RAG scanning. Nature 590, 338–343 (2021).

46. B. Nabet et al., Rapid and direct control of target protein levels with VHL-recruiting dTAG molecules. Nat Commun 11, 4687 (2020).

47. G. D. Victora, M. C. Nussenzweig, Germinal Centers. Annu Rev Immunol 40, 413–442 (2022).

48. A. Peters, U. Storb, Somatic Hypermutation of Immunoglobulin Genes Is Linked to Transcription Initiation. Immunity 4, 57–65 (1996).

49. R. Pavri et al., Activation-induced cytidine deaminase targets DNA at sites of RNA polymerase II stalling by interaction with Spt5. Cell 143, 122–133 (2010).

50. Y. Zhang et al., Transcriptionally active HERV-H retrotransposons demarcate topologically associating domains in human pluripotent stem cells. Nat Genet 51, 1380–1388 (2019).

51. S. P. Methot et al., A licensing step links AID to transcription elongation for mutagenesis in B cells. Nat Commun 9, 1248 (2018).

52. X. Zhang, H. S. Yoon, A. M. Chapdelaine-Williams, N. Kyritsis, F. W. Alt, Physiological role of the 3’IgH CBEs super-anchor in antibody class switching. Proc Natl Acad Sci U S A 118 (2021).

53. A. M. Patenaude et al., Active nuclear import and cytoplasmic retention of activation-induced deaminase. Nat Struct Mol Biol 16, 517–527 (2009).

54. Q. Le, N. Maizels, Activation-induced deaminase (AID) localizes to the nucleus in brief pulses. PLoS Genet 15, e1007968 (2019).

55. B. Laffleur et al., Noncoding RNA processing by DIS3 regulates chromosomal architecture and somatic hypermutation in B cells. Nat Genet 53, 230–242 (2021).

56. V. Delgado-Benito et al., The Chromatin Reader ZMYND8 Regulates Igh Enhancers to Promote Immunoglobulin Class Switch Recombination. Mol Cell 72, 636–649 e638 (2018).

57. J. Hu et al., Detecting DNA double-stranded breaks in mammalian genomes by linear amplification-mediated high-throughput genome-wide translocation sequencing. Nat Protoc 11, 853–871 (2016).

58. D. B. Mahat et al., Base-pair-resolution genome-wide mapping of active RNA polymerases using precision nuclear run-on (PRO-seq). Nat Protoc 11, 1455–1476 (2016).

