## Supplemental data for "Human antibody heavy chain variable region exon interacts with 3’ regulatory region to form a stable somatic hypermutation center"

**This PDF file includes:**

Figures S1 to S8
Tables S1 to S3

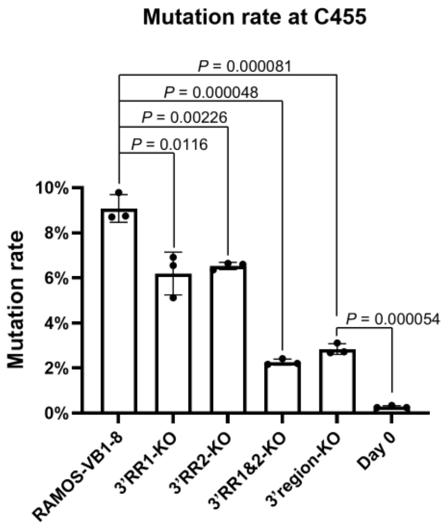


**Fig. S1.** Mutation frequency at the C455 nucleotide position of the VB1-8 exon in RAMOS-VB1-8, 3′RR1-KO, 3′RR2-KO, 3′RR1&2-KO, and 3′region-KO cells after 10 days of doxycycline treatment, as determined by SHM-Rep-Seq. The day 0 sample from untreated RAMOS-VB1-8 cells served as a negative control. Data are shown as mean ± SD from three independent repeats.

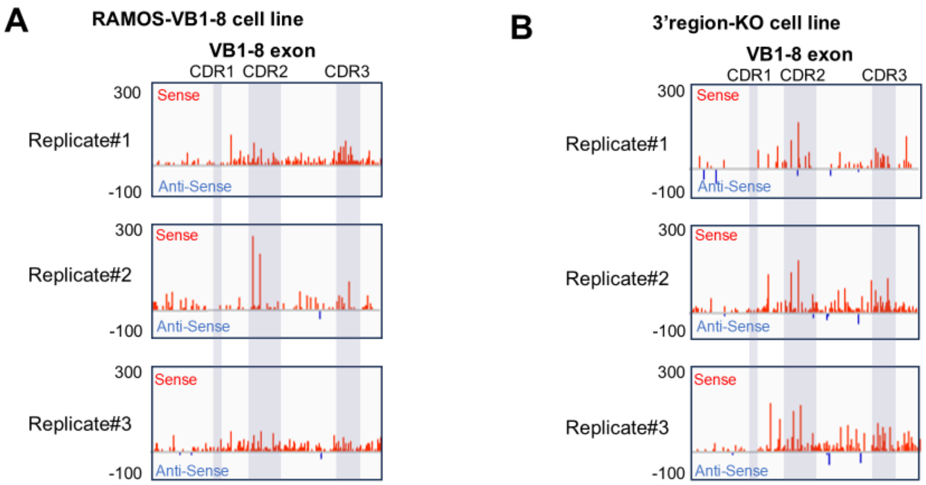


**Fig. S2.** Transcription levels of the VB1-8 exon in RAMOS-VB1-8 and 3′region-KO cells across independent replicates. (A,B) PRO-seq reads across the VB1-8 exon in RAMOS-VB1-8 cells (A) and 3′region-KO cells (B).

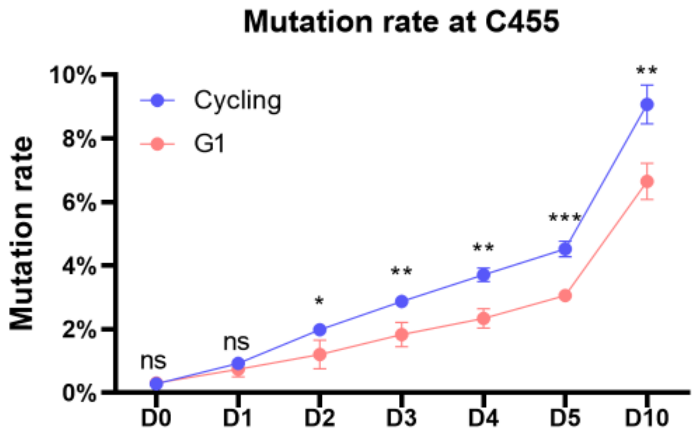


**Fig. S3.** Mutation rate at the C455 nucleotide position of the VB1-8 exon in cycling RAMOS-VB1-8 cells and G1-arrested RAMOS-VB1-8-Bcl2 cells, as determined by SHM-Rep-Seq. Data represent the mean ± SD of three independent repeats. * $P < 0.05$, ** $P < 0.01$, *** $P < 0.002$; ns, not significant.

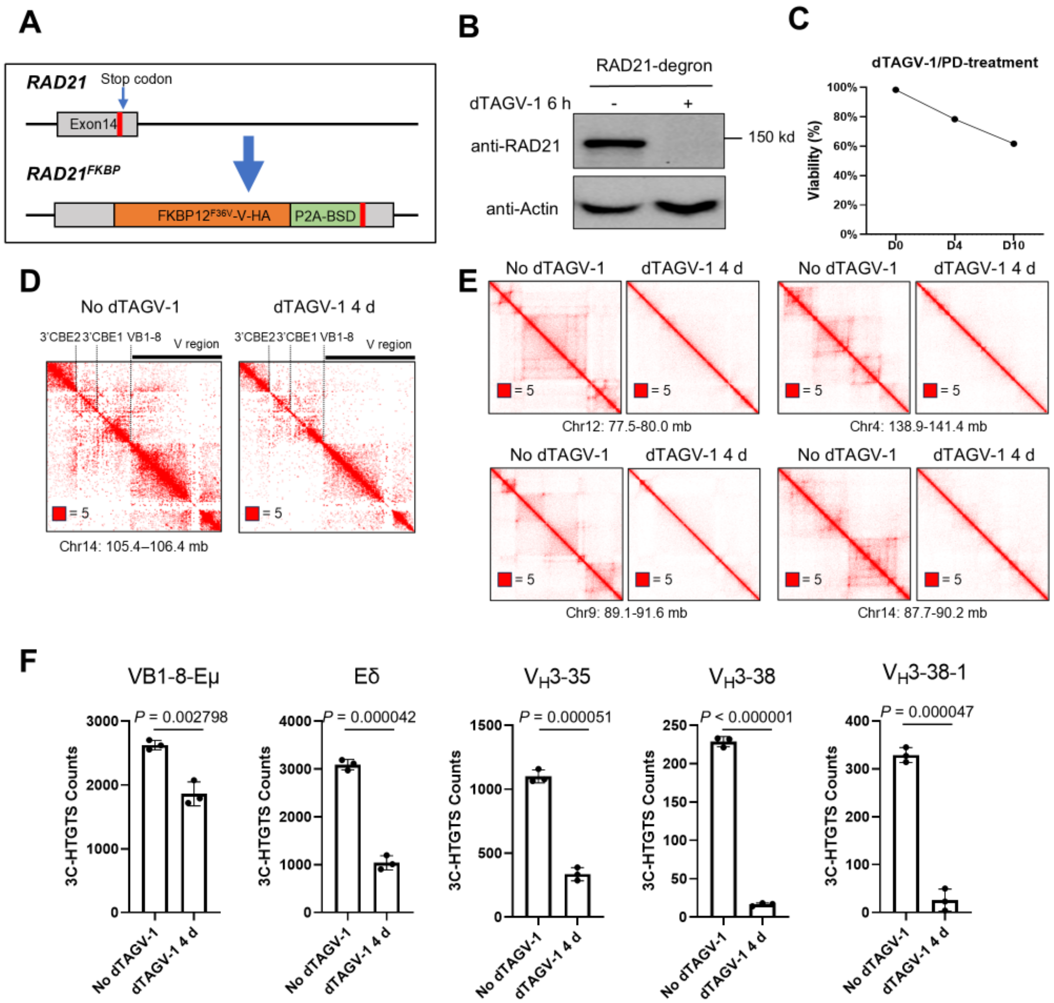


**Figure S4.** Validation of RAD21 depletion and its effects on chromatin architecture and transcription at the *IGH* locus. (A) Schematic of the strategy used to generate RAD21-degron cells. (B) Western blot analysis of RAD21 protein levels in RAD21-degron cells treated with or without dTAGV-1 for 6 hours. Raw data is shown in Figure S8B. (C) Cell viability of RAD21-degron cells following 10 days of dTAGV-1 treatment. Data represent the mean ± SD of three independent repeats. (D) Hi-C interaction maps of the *IGH* locus in RAD21-degron cells treated with or without dTAGV-1 for 4 days. (E) Hi-C interaction maps of four non-*IGH* loci in RAD21-degron cells treated with or without dTAGV-1 for 4 days. (F) Normalized 3C-HTGTS contact counts for each indicated 5-kb region from 3′RR-baited 3C-HTGTS libraries in RAD21-degron cells treated with or without dTAGV-1. Data are shown as mean ± SD from three independent repeats.

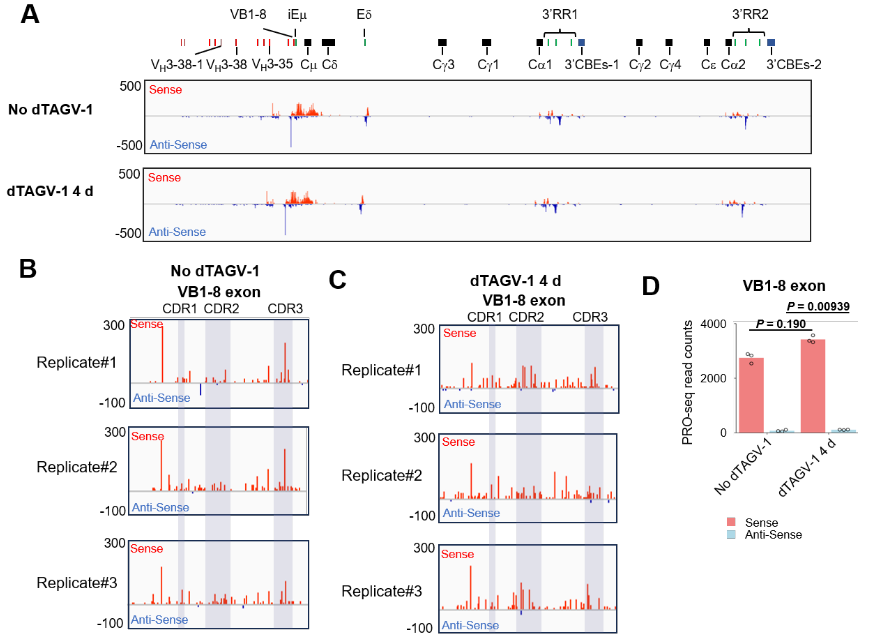


**Figure S5.** Transcription levels of the VB1-8 exon in RAD21-degron cells with and without dTAGV-1 treatment of independent replicates. (A-C) Normalized PRO-seq read counts across the *IGH* constant region (A) and the VB1-8 exon (B-C) in RAD21-degron cells treated with or without dTAGV-1 for 4 days. (D) Quantification of normalized PRO-seq read counts across the VB1-8 exon in RAD21-degron cells treated with or without dTAGV-1 for 4 days. Statistical significance was assessed using a two-tailed unpaired Student's t-test. Data represent the mean of three independent repeats.

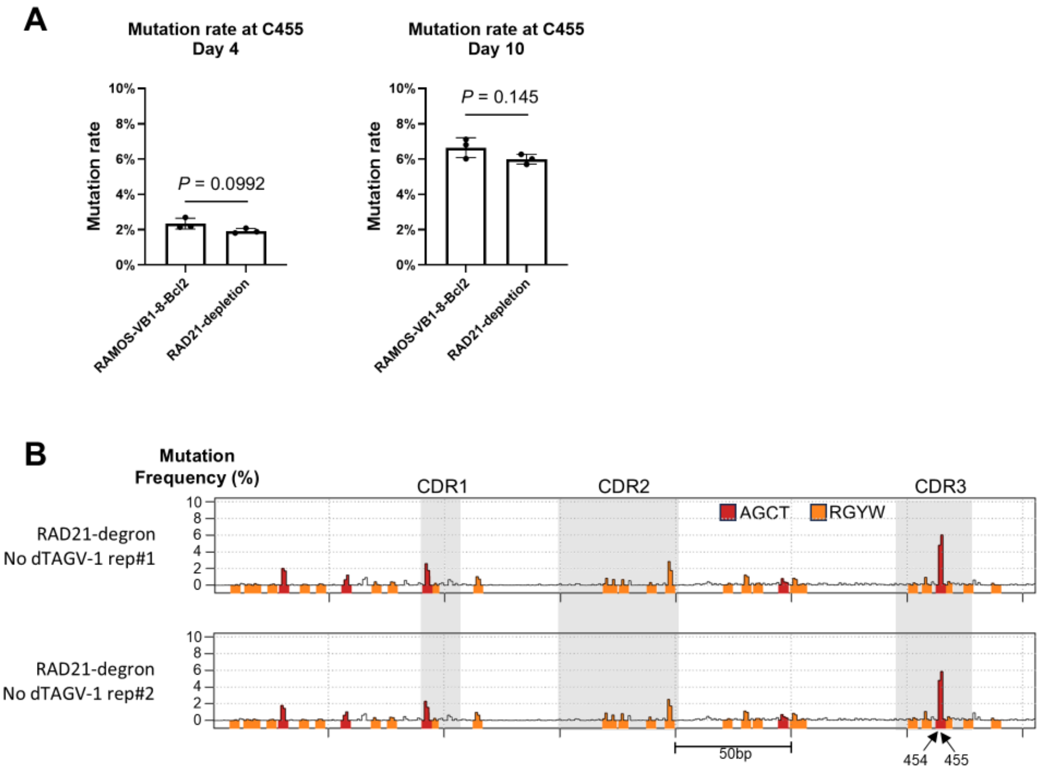


**Figure S6.** VB1-8 mutation efficiency in RAD21-degron cells. (A) Mutation frequency at the C455 site in RAMOS-VB1-8-Bcl2 and RAD21-depleted cells after 4 or 10 days of doxycycline treatment, as determined by SHM-Rep-Seq. Data represent the mean ± SD from three independent repeats. (B) SHM profiles across the VB1-8 exon in RAD21-degron cells following 10 days of doxycycline-induced AID expression without dTAGV-1 treatment, as determined by SHM-Rep-Seq. Data are shown from two independent replicates.

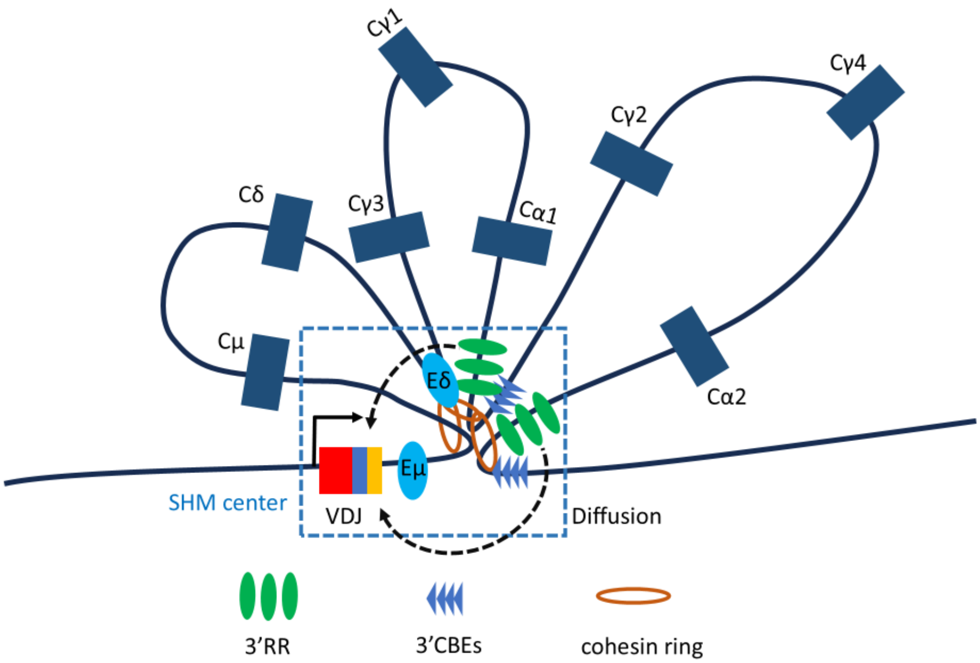


**Figure S7**. This figure depicts one model for the structure of the proposed SHM-C, based on the data described in the manuscript. We propose that in GC B cells that cohesin is loaded within each set of $C_H$ exons sequences, potentially within the 3'RRs and/or the upstream Eμ/Sμ region. Extrusion through the upstream $C_H$ exons would be unimpeded but then dynamically impeded by the transcribed iEμ/Sμ sequences upstream and the transcribed 3'RR1/3'CBE1 downstream to form a dynamic upstream loop that juxtaposes the VDJ and 3'RR1/3'CBE1 in one domain of the SHM center. Extrusion from cohesin loaded in the downstream $C_H$/3'RR sequences would be dynamically impeded the transcribed 3'RR2/3'CBE2 downstream and by the 3'RR1/3'CBE1 upstream to form a dynamic downstream loop that juxtaposes 3'RR2 and 3'RR1 to establish the second domain of the SHM center. This model would position both 3'RRs within short-range diffusion distance from the V(D)J exon and thereby able to support SHM activity. Both the upstream-based and downstream-based downstream domains could form an active 3'RR1-based or 3'RR2-based SHM-C in the absence of the other in the context of this model, explaining our finding that one or the other 3'RR can support SHM. Please see text or the Supplementary Discussion of reference 13 for more details.

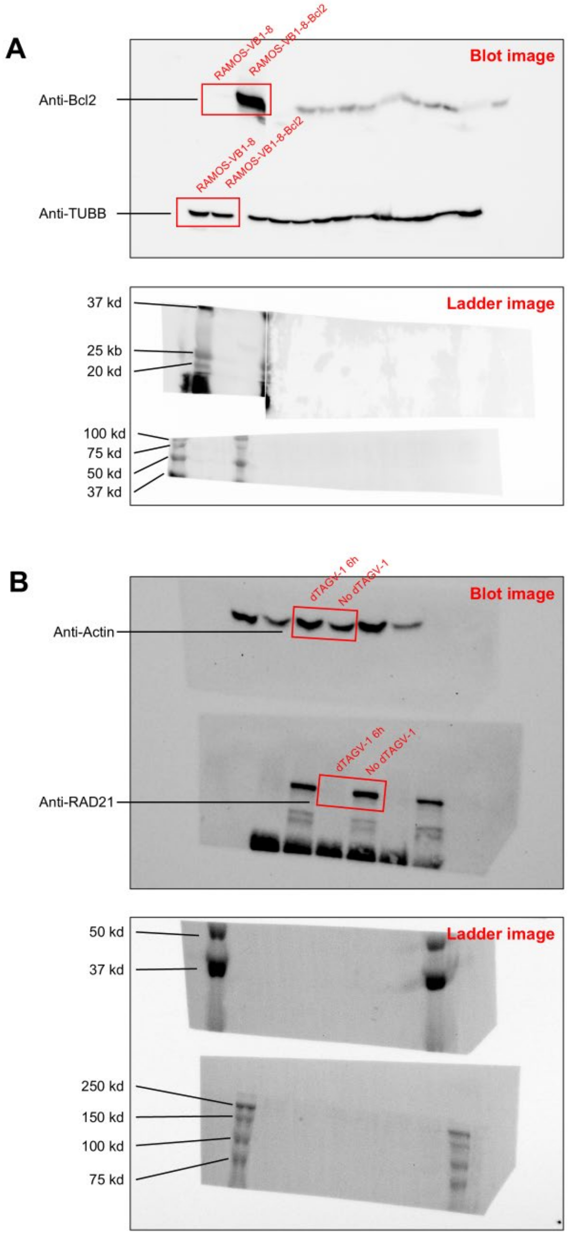


**Figure S8**. Unprocessed western blot analysis. (A) Bcl2 western blot for RAMOS-VB1-8 and RAMOS-VB1-8-Bcl2 cells. Top, blot image; bottom, ladder image. Red boxes indicate the regions presented in Figure 4D. The blot was generated from the same membrane, which was cut prior to probing with different antibodies. (B) RAD21 western blot for RAD21-degron cells treated with or without dTAGV-1 for 6 hours. Top, blot image; bottom, ladder image. Red boxes indicate the regions presented in Figure S4B. The blot was generated from the same membrane, which was cut prior to probing with different antibodies.

Tables

**Table S1.** List of sgRNAs

| Name | Sequence | Purpose |
|---|---|---|
| sgRNA-Ramos VDJ 5'-2 | CGGCGTGTAGACATCTACGT | Targeting the endogenous VDJ locus upstream, for VB1-8 insertion |
| 3gRNA-vb18p-new1 | CAATGCTCAGGAAACCCCAC | Targeting the endogenous VDJ locus downstream, for VB1-8 insertion |
| RRCBE-sg4 | CCGACCCGCCCTTACTCACG | Targeting the upstream of 3'RR-2 |
| RRCBE-sg5 | GTGGAGATATGCGGAAGTGT | Targeting the downstream of 3'RR-2 |
| RRCBE-sg7 | GCCTTCCAGGTGCAATAAAG | Targeting the upstream of 3'RR-1 |
| RRCBE-sg2 | GTGAAGGTGTGAGAGTGTGA | Targeting the downstream of 3'RR-1 |
| grna-rr1cbe1 | CTGCAGGTCACTTACTCATG | Targeting the upstream of 3' region |
| grna-rr2cbe2 | GACGTCCTGACCACCCTAAT | Targeting the downstream of 3' region |
| AAVS1 sgRNA-4 | GGGGCCACTAGGGACAGGAT | Targeting the AAVS1 locus, for Bcl2 expression cassette insertion |
| sgRAD21 | CCAAGGTTCCATATTATATA | Targeting the RAD21 locus, for FKBP tag insertion |

**Table S2.** List of oligos.

| Name | Sequence | Purpose |
|---|---|---|
| bio-vb18-F3 | ACATATATATGGGTGACAATGACATC | bio primer for VB1-8 bait for SHM-Rep-seq and 3C-HTGTS |
| red-vb18 | AATGACATCCACTTTGCCTTTCTCT | red primer for VB1-8 bait for SHM-Rep-seq |
| VB18-red-3C | GGCCACACTGACTGTAGACAAAC | red primer for VB1-8 bait for 3C-HTGTS |
| bio-HS12-loca1-F1 | GTCCTGGTCCCAAAGATGGC | bio primer for 3'RR bait for 3C-HTGTS |
| red-HS12-loca1 | AAGCTATTTCTGGAAACAGCCTCAG | red primer for 3'RR bait for 3C-HTGTS |

**Table S3.** List of antibodies.

| Name | Source | Purpose and dilution ratio |
|---|---|---|
| Anti-Bcl2 | Abcam, # ab182858 | Western blot, 1:2000 |
| Anti-TUBB | Abcam, # ab78078 | Western blot, 1:1000 |
| Anti-RAD21 | Abcam, # ab154769 | CUT&RUN, 1:100 |
| Anti-RAD21 | Abcam, # ab992 | Western blot, 1:1000 |
| Anti-Actin | CST, # 5125 | Western blot, 1:4000 |
| Anti-AID | Home made | CUT&RUN, 1:100 |
